# Distinct growth regimes govern crowding in foveal and extrafoveal vision

**DOI:** 10.1101/2025.09.17.673105

**Authors:** Ashley M. Clark, Krishnamachari S. Prahalad, Martina Poletti

## Abstract

Visual crowding—where object recognition is impaired by nearby stimuli—is a well-documented phenomenon in the visual periphery, thought to reflect fundamental processes of grouping and segmentation. A hallmark of crowding is its spatial extent increasing linearly with eccentricity, as described by Bouma’s law, reflecting retinal convergence and cortical magnification. Although crowding is typically studied in the periphery—and Bouma’s law predicts no crowding at the center of gaze—it occurs even foveally. Within the foveola, the central 1 deg of the visual field, characterized by peak visual acuity and largely one-to-one connectivity between photoreceptors and ganglion cells, the Bouma law breaks and the extent of crowding is expected to remain constant with eccentricity. Yet, whether crowding varies at this fine spatial scale remains unknown. To investigate this, we combined high-resolution eye tracking with a gaze-contingent display system to precisely localize gaze and measure crowding thresholds at multiple foveolar eccentricities. Our results reveal that crowding does increase linearly within the foveola, but at a rate significantly slower—approximately 3.5 times-than in the extrafovea. By showing that even within the highest-acuity region of the visual field, spatial integration zones are not fixed but change with eccentricity these findings challenge the view of the central fovea as a spatially uniform processing zone and demonstrate that crowding at this scale follows a distinct regime.

**Significance:** When we view objects in clutter, recognition declines—a phenomenon known as crowding. This effect increases with distance from the center of gaze, following Bouma’s law, which predicts little or no crowding at the fovea’s center. As this law breaks at the center of gaze, it has been hypothesized that around this region, crowding remains constant with eccentricity. Using high-resolution eyetracking, we show that even within the central 0.4*°* of the visual field, crowding increases steadily with minute stimulus offsets from the point of fixation. This growth is much slower than in peripheral vision, revealing a distinct crowding regime in the foveola, likely reflecting cortical and retinal differences. Our findings offer new insights into the fine-scale organization of spatial vision.

## Introduction

Imagine you are driving during rush hour on a crowded urban road. In addition to vehicles, there are pedestrians and cyclists moving in and out of traffic. The constant flow of people and bikes creates a complex visual scene, making it challenging to identify potential hazards or predict their movements. This difficulty arises in part from a phenomenon known as visual crowding; recognizing an object becomes harder when it is surrounded by similar stimuli, generally referred to as flankers (see [1–7] for a review on crowding). As the distance between target and flankers increases, the detrimental effect of crowding is minimized. The smallest distance between the target and flanker at which performance drops by a given amount from the asymptotic performance level is referred to as critical spacing, which defines the spatial extent of the visual crowding effect. However, the extent of crowding is not uniform across the visual field; it depends on the target eccentricity from the center of gaze, a principle known as Bouma’s law [8, 9]. According to this law, the extent of visual crowding grows linearly with eccentricity, [8, 9]; as the target moves farther away from the center of gaze, the distance between the flankers and the target needs to be increased to maintain similar performance levels across eccentricities.

Visual crowding has been extensively studied in the visual periphery where its detrimental effects are most pronounced [1, 9]. Meanwhile, crowding at the foveal level is thought to have minimal impact on object recognition, as visual acuity is considered the main bottleneck at this scale [1, 2]. Although the existence of crowding in the central fovea has been debated over the past sixty years [2, 10, 11], there is now mounting evidence showing that crowding also influences vision also at the very center of gaze where acuity is highest [12–17]. Further, the effects of foveal crowding have been reported even when stimuli are presented under diffraction limited conditions, correcting for optical aberrations, which under normal viewing conditions may act as a confounding factor for crowding at this scale [18].

The 1*^◦^* central fovea, or foveola, is anatomically different from the rest of the retina. It is situated in a pit, free from rods and capillaries, and contains densely packed cone photoreceptors, thus facilitating fine spatial vision [19]. Traditionally foveolar vision is studied as a unit, and there is limited understanding of how it varies at a sub-degree scale. Visual detection thresholds were found to be relatively uniform within the central foveal region ([20], but see [21]). On the other hand, recent research has shown that the ability to discriminate fine details is not uniform and it drops already 15’ away from the preferred locus of fixation (PLF) [22, 23]. This raises the question of whether the effects of crowding are the same across the central 1-deg foveal region or if they are modulated by changes in eccentricity even at this finer scale.

Visual processing in the fovea and periphery is shaped by striking differences in retinal structure and cortical representation. In the fovea, the unique one-to-one mapping between photoreceptors, bipolar and retinal ganglion cells (RGCs), allows for high-fidelity spatial resolution and color discrimination [19, 24]. This high-resolution processing is supported by private-line wiring, where each cone photoreceptor connects to dedicated ON and OFF bipolar cells, which in turn project to distinct midget RGCs, ensuring precise signal transmission and minimal convergence [25–27]. In contrast, the more peripheral retina exhibits significant convergence, where multiple photoreceptors converge onto a single RGC, resulting in spatial pooling that prioritizes sensitivity over fine detail [24]. These retinal differences manifest in the primary visual cortex (V1) as a disproportionate allocation of cortical area to foveal processing, described by the cortical magnification factor (M) [28, 29]. This measure quantifies how much cortical tissue is devoted to different regions of the visual field, with the fovea receiving vastly more resources compared to the periphery [30, 31]. Such cortical organization allows for precise spatial discrimination in foveal vision [32]. In addition to its impact on visual resolution, it has been proposed that RGC convergence and cortical magnification influence visual crowding, with studies suggesting that the spatial extent of crowding may depend on the size of RGC receptive fields [33] and cortical magnification [5, 34, 35] along the visual pathway. It was unclear until recently whether cortical magnification is uniform within the foveola, however, recent evidence from non-human primates indicates that a cortical magnification gradient is present even within the foveola [36], and this gradient can potentially affect foveal crowding. Hence, important questions about the specific mechanisms underlying crowding within the fovea remain open.

The discussion on foveal crowding also raises questions about potential differences in the effects and mechanisms of crowding in the 1-deg central fovea compared to extrafoveally. These variations may arise from distinctions in neural architecture at the cortical and/or retinal level and in different functional demands between these retinal regions. The central fovea, optimized for high acuity, contrasts with the periphery, emphasizing motion sensitivity and broader visual awareness (see [2] for a review on this). The structural and functional differences between the foveola and the peripheral visual field suggest that the factors influencing visual crowding may vary between these two regions. Three possible scenarios for how crowding might be characterized within the foveola can be outlined (see Figure 1*a*): (1) as with Bouma’s law breaks at the center of gaze, it is conceivable that the impact of crowding does not change with eccentricity within the foveola, (2) Alternatively, the region of uniform crowding effects may only be limited to the center of gaze and to a small surrounding zone within the foveola. However, beyond this zone crowding extent may increase with eccentricity within the foveola following the same Bouma’s law as extrafoveally [37]; (3) A third possibility is that the increase of crowding extent with eccentricity is still present in the foveola, but it follows a different regime, possibly reflecting differences in cortical magnification and/or physiological changes between this region and the rest of the retina. In particular, results from Rossi and Roorda (2010) suggest that the private-line connection between cones and Retinal Ganglion Cells (RGCs) is limited to the central 1 deg diam region around the preferred retinal locus. A sudden increase in the rate of convergence, which is then reflected in cortical magnification in V1 and beyond, may lead to two distinct crowding regimes in the foveola and extrafoveally.

**Figure 1:**
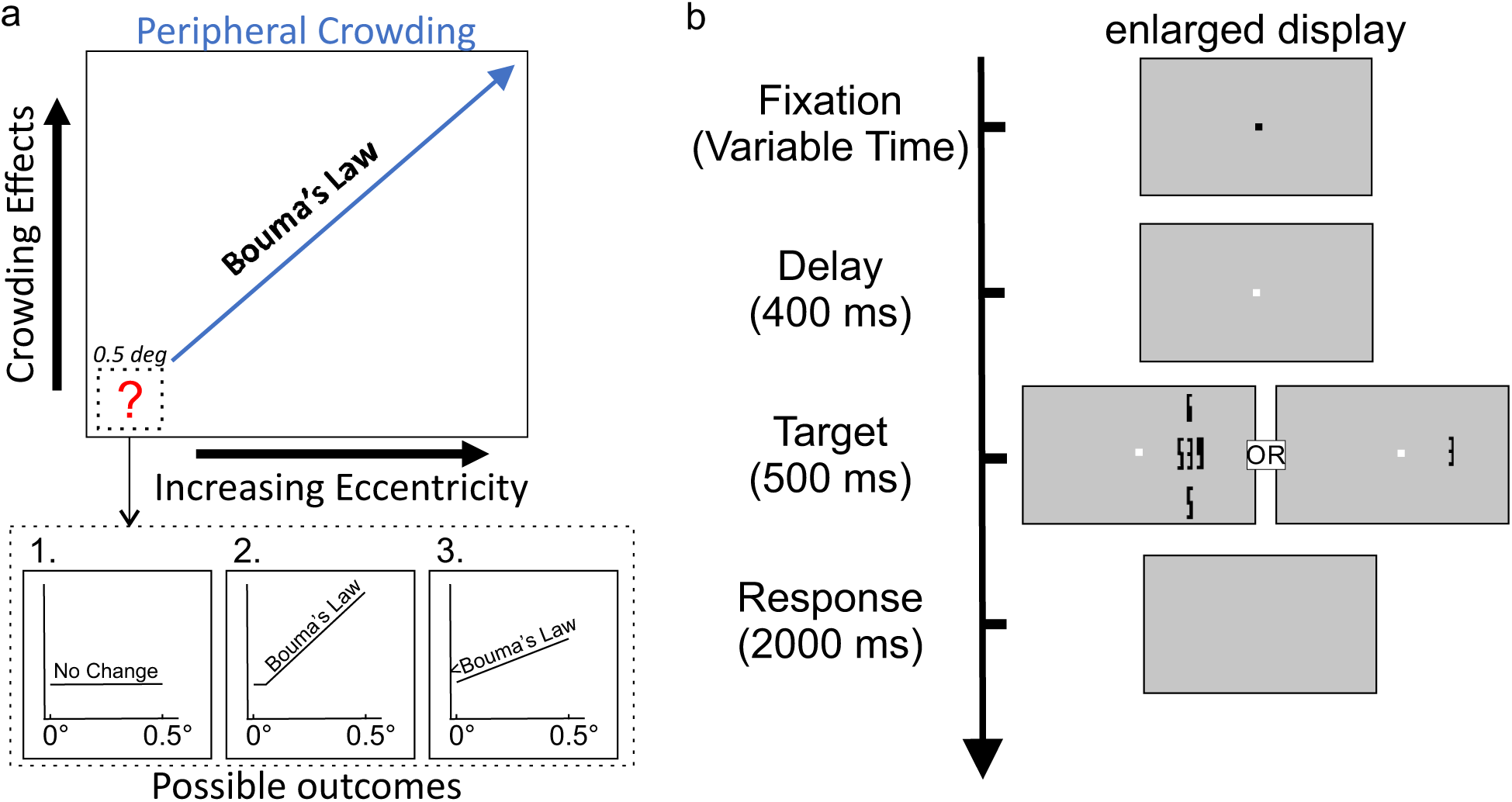
Crowding in the Foveola. (a) An illustration of how crowding magnitude increases with eccentricity following the Bouma’s Law is well known to increase crowding effects extrafoveoally. Yet it is unknown if and how the magnitude of crowding changed within the central 1-degree diameter foveola. *Bottom*, three predictions on how the relationship between crowding and eccentricity may unfold within the foveola: 1.Bouma’s law breaks in this region where crowding is still observed and the effects of crowding do not change across the foveola; 2. The region of uniform crowding effects is limited to a small zone around the center of gaze and beyond this limit crowding follows the same Bouma’s law it does extrafoveally; 3. crowding increases with eccentricity within the foveola, but with a slower rate compared to extrafoveally. (b) Experimental Paradigm. Subjects were instructed to identify high-acuity stimuli presented at the center of the display for 500 ms. Stimuli appeared after a brief period of fixation and a 400 ms period in which the display was blank, in the 0’ eccentricity condition, to avoid possible after effects of the central fixation dot. Stimuli, digits in Pelli font [12], varied in size from 0.3’ to 4’ and in eccentricity presented at either 0, 10, 15, or 25-arcmin eccentricity. Observers were asked to identify the central digit (the target) among 4 possible choices.

The foveola is often treated as a functionally uniform region, in part because retinal image quality remains relatively stable across this area [39–42], and because optical aberrations limit visual resolution to well below the Nyquist sampling limit of the foveal cone mosaic [38]. Further, spotlight sensitivity within the foveola has been reported to be uniform even under diffraction limited viewing [20]. Yet, cone density changes sharply within the foveola [20, 24, 43], fine spatial vision declines with increasing eccentricity within this region [22, 23], and contrast detection has been shown to vary within the foveola under normal viewing conditions in the presence of noise [23]. It therefore remains unclear whether the mechanisms governing higher-order visual functions, such as grouping and segmentation, operate uniformly across the foveola or if they are modulated with eccentricity, as it happens in the extrafoveal visual field (Fig. 1*a*). To address this question, we examined the phenomenon of visual crowding, the direct consequence of grouping and segmentation processes [4], across the foveola. Previous studies have shown that the central fovea is not immune to crowding [12, 14–17, 44, 45], and that crowding persists even under diffraction-limited conditions achieved with adaptive optics [18]. Here we investigate whether the extent of crowding varies within the foveola under natural viewing conditions—that is, in the presence of physiological higher-order aberrations and fixational eye movements.

Fixational eye movements, though critical for fine vision (see review [21]), pose technical challenges for assessing acuity and crowding near the center of gaze. Eye motion during fixation [17, 46–48] and spatial imprecision in video-based eye trackers [49–51] make it difficult to deliver stimuli just a few arcminutes from the foveal center. To overcome this, we used high-precision eye tracking with real-time, gaze-contingent stimulus delivery [52, 53]. While retinal stabilization might circumvent these challenges, it imposes highly unnatural viewing conditions and it leads to a drop in contrast sensitivity [54]. In the current study, we allowed for natural fixational eye movements to occur while simultaneously ensuring that stimulation remained confined to the desired eccentricity.

## Results

To measure the effects of crowding across the foveola, visual acuity was first assessed using a 4AFC visual discrimination task. Stimuli consisted of digits in Pelli font, a font specifically designed for testing foveal crowding [12] (Figure 1*b*). The width of the stimuli ranged from 0.4 to 5 arcminutes, with the height scaled by a factor of 5 (*i.e.*, an aspect ratio of 1:5, width:height, as in [12]). Stimulus size was adjusted using an adaptive staircase procedure to determine the threshold acuity at each eccentricity [55]. To assess crowding, critical spacing thresholds were measured by surrounding the target with four flanker digits. Flankers were positioned along the horizontal and vertical axes, with center-to-center spacing set to 1.4 times the width and height of the target along each axis, respectively. As a result, in the crowded condition, center-to-center spacing scaled proportionally with stimulus size. Because crowding in this font primarily depends on horizontal spacing, crowding effects are referred to horizontal center-to-center spacing. This design allowed us to quantify both acuity and crowding simultaneously.

To ensure precise eye movement measurements and to limit visual stimulation around the desired foveal eccentricity, we employed a high-precision eyetracker [52] coupled with a state-of-the-art custom-made gaze-contingent display system [53]. The combination of these systems allowed for more accurate gaze localization during the experimental tasks [21]. Figure 2 shows the gaze distribution maps across all tested conditions. To reflect a more natural viewing condition while still ensuring accurate stimulus placement, we allowed for the temporal modulations introduced by fixational eye movements but excluded trials with excessive deviation from the central marker and trials with microsaccades; only trials in which gaze position remained within 30 arcminutes of the center of the display were included in the analysis. As shown in Figure 2, in the selected trials, 85.33% ± 7.46% of gaze positions remained within a ± 10 arcminute region around the center of the display. The average euclidean distance from the display center was 1.03 ± 0.49 arcminutes in the unflanked condition and 1.07 ± 0.50 arcminutes in the crowded condition. A two-way repeated-measures ANOVA with within-subject factors of eccentricity and crowding revealed no significant main effect of crowding on fixation accuracy (F(3,5) = 0.0008, p = 0.98, *η*^2^ *<* 0.001), no significant main effect of eccentricity (F(3,15) = 0.71, p = 0.56, *η*^2^ = 0.021), and no interaction between eccentricity and crowding (F(3,15) = 1.39, p = 0.29, *η*^2^ = 0.0065). These results indicate that fixation accuracy was comparable in both conditions and across eccentricities.

**Figure 2:**
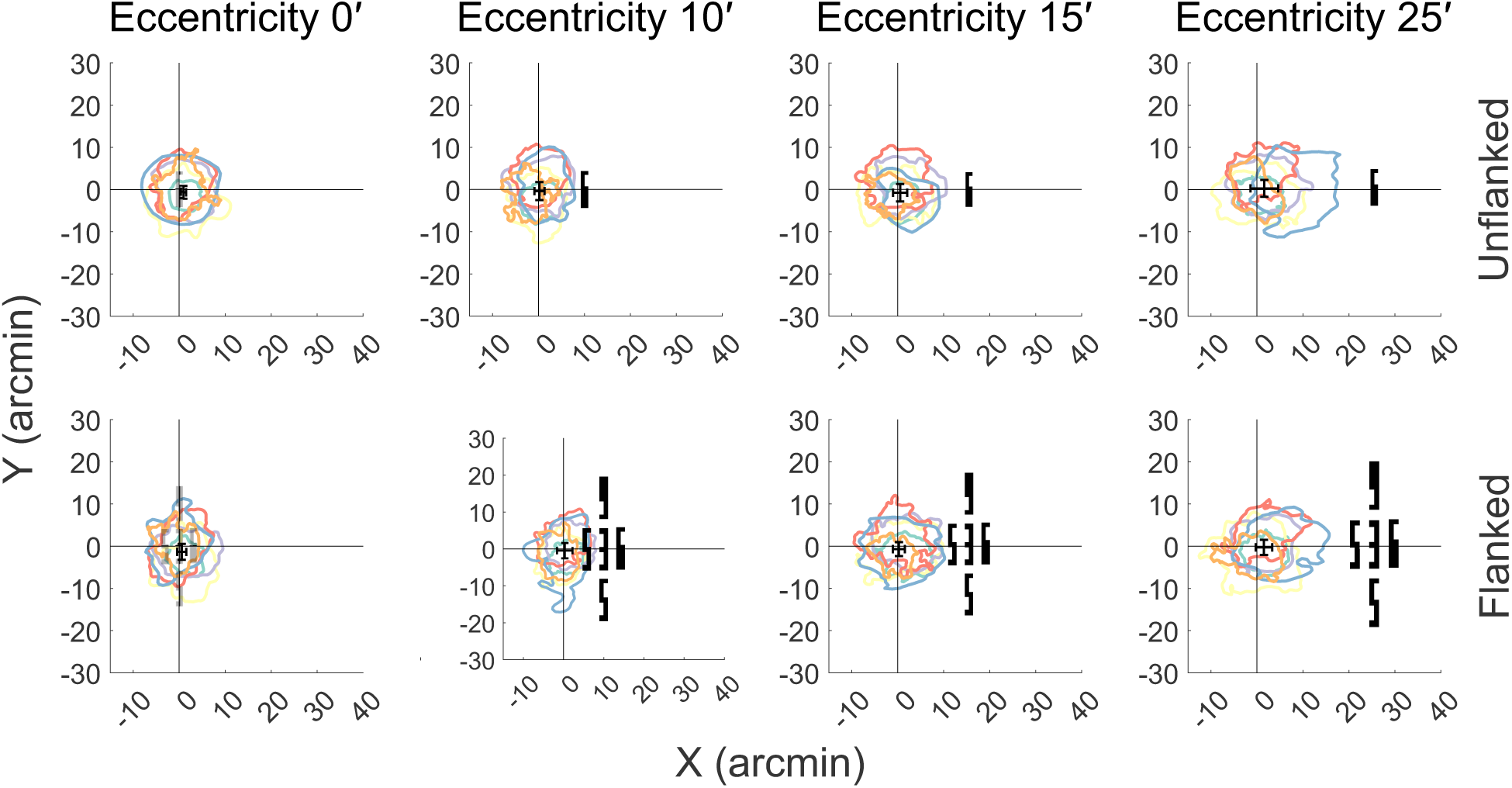
Gaze position during stimulus presentation. 2D contour plots highlight the area encompassing 68% of the gaze positions during the stimulus presentation. Gaze distributions for each individual are shown in separate colors for the isolated/unflanked (top row) and flanked (bottom row) conditions during stimulus presentation interval. An example stimulus is overlaid in each panel (except at 0’ eccentricity) to illustrate the spatial position of the target relative to the center of the display. The stimulus is scaled based on the average threshold size across subjects in each condition. Individual columns represent the distributions for the different eccentricities tested. When stimuli were presented away from the PLF or *>* 0’ eccentricity, subjects maintained their gaze on the fixation marker, ensuring that retinal stimulation occurred at the desired eccentricity.

When stimuli were presented in isolation (unflanked) at the PLF, the average stimulus width required to reach threshold performance levels was 1.62’ ± 0.21’, approximately equivalent to 20/16 Snellen acuity. Acuity thresholds in the unflanked condition (isolated stimuli) remained relatively constant across the eccentricities (repeated measures ANOVA: F(3,15) = 1.05, p = 0.399, *η*^2^ = 0.0007), whereas thresholds in the presence of flankers (crowded stimuli) significantly increased with eccentricity (repeated-measures ANOVA: F(3,15) = 9.56, p *<* 0.001, *η*^2^ = 0.012), indicating a systematic degradation in visual acuity under crowded conditions (red circles, Figure 3*a*) compared to isolated stimulus presentation (blue circles, Figure 3*a*). As previously shown [17], at the center of gaze, there was an increase of 29.07%± 9.42% increase in acuity thresholds (approximately equivalent to 20/20 Snellen acuity) relative to the unflanked condition (two-tailed paired t-test, t(5) = 8.40, p =0.0004, *BF*_10_= 82.69, Cohen’s D= 1.93).

**Figure 3:**
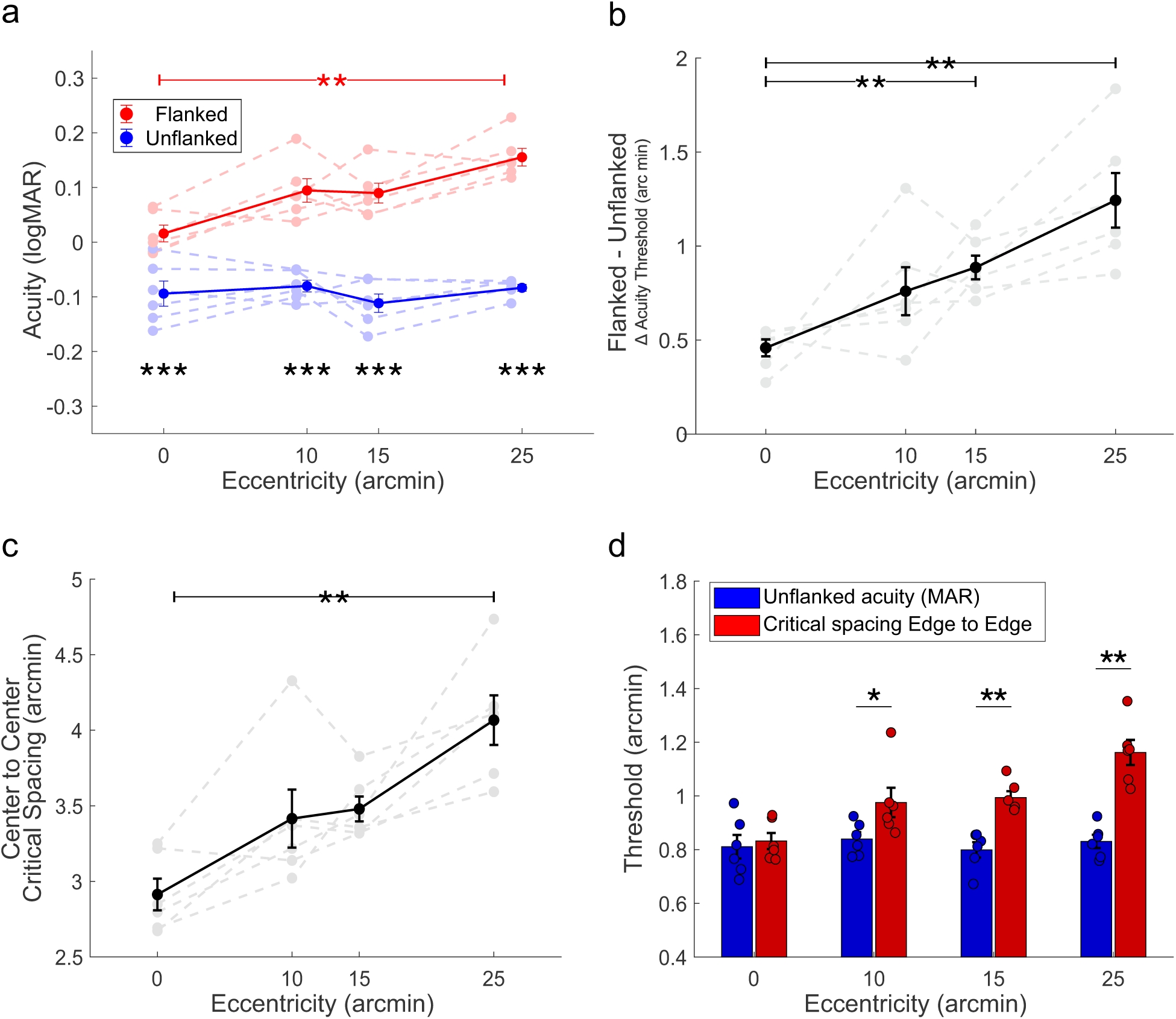
Effects of crowding on visual acuity. (a) Crowded acuity declined with increasing eccentricity (red circles) and remained worse than unflanked acuity (blue circles) at each eccentricity(repeated-measures ANOVA: *p* = 0.005 for eccentricity, *p <* 0.001 for flanker condition, *p* = 0.001 for interaction). (b) Difference in acuity thresholds between the crowded and unflanked condition for the different eccentricities tested. The difference increased with the increase in stimulus eccentricity (repeated measures ANOVA: *p <* 0.001). (c) Critical spacing thresholds, represented as the center-to-center separation between the target and the flanker for crowded stimuli, as a function of eccentricity. Similarly, we observed an increase in critical spacing thresholds with eccentricity (repeated measures ANOVA: *p <* 0.001). d) Threshold estimates for unflanked acuity and critical spacing estimates as edge-to-edge spacing for flanked stimuli as a function of eccentricity. Error bars indicate s.e.m.. Asterisks indicate statistically significant post-hoc pairwise comparisons performed using the Tukey-Kramer procedure across eccentricities in panels (a–c) and significant differences between critical spacing and unflanked MAR at each eccentricity in panel (d) (^∗^*p <* 0.05, ^∗∗^*p <* 0.01).

A two-way repeated-measures ANOVA with eccentricity and flanker condition as within-subject factors revealed significant main effects of eccentricity (*F* (3,15) = 6.43, *p* = 0.0052, *η*^2^ = 0.068) and flanker condition (*F* (1,5) = 328.42, *p <* 0.001, *η*^2^ = 0.752), as well as a significant interaction (*F* (3,15) = 9.10, *p* = 0.0011, *η*^2^ = 0.051). Post hoc comparisons showed that flanked thresholds were elevated relative to unflanked thresholds at each eccentricity tested (all eccentricities *p <* 0.001). Within the flanked condition, 25’ thresholds were significantly higher than 0’ (p = 0.005), while other flanked pairwise comparisons and all unflanked comparisons across eccentricities were not significant (p = 0.057 and p = 0.105, respectively). Flanked visual acuity thresholds increased by 45.4% at 10’, 56.3% at 15’, and 76.1% at 25’ relative to unflanked their respective unflanked conditions. Further, we calculated the difference in visual acuity thresholds (in arcminutes) between the flanked and unflanked conditions at each tested eccentricity. When plotting the threshold difference as a function of eccentricity (Figure 3*b*), all subjects showed consistently positive differences, indicating stronger crowding effects at greater eccentricities. A repeated-measures ANOVA confirmed that crowding strength increased with eccentricity (F(3,15) = 9.42, *p <* 0.001, *η*^2^ = 0.085). Post hoc comparisons revealed that crowding at 15 and 25 arcminutes was significantly greater than at 0 arcminutes (p = 0.004 and p = 0.006, respectively), while the difference between 0 and 10 arcminutes did not reach significance (p = 0.171).

To complement acuity based measures, we examined critical spacing, the center-to-center distance between the target and flanker at the acuity threshold, providing a more direct measure of the spatial extent of crowding. We observed a systematic increase in the critical spacing measures with increasing stimulus eccentricity (repeated measures ANOVA: F(3,15) = 9.56, *p <* 0.001, *η*^2^ = 0.012). Post hoc comparisons revealed that critical spacing at 25’ was significantly greater than at 0’ (p = 0.007), while the differences between 0’ and 10’ and between 0’ and 15’ did not reach significance (p = 0.067 and p = 0.125 respectively) (figure 3*c*). In other words, as eccentricity increased within the foveola, critical spacing (center-to-center distance) increased from 2.91’± 0.1’ at the PLF to 4.02’ ± 0.16’ at 25’ eccentricity, indicating that flankers needed to be placed farther from the target to maintain similar performance levels.

Beyond measuring the spatial extent of crowding, we compared these values with resolution limits derived from unflanked stimulus thresholds. Specifically, we compared the minimum angle of resolution (MAR), which reflects the limit of visual acuity, to the crowding limit, this time defined as the edge-to-edge (E-E), instead of the center-to-center, critical spacing threshold. This allowed us to compare the acuity limits with the crowding limits across the tested eccentricities within the foveola (see supplemental figure S1). The minimum angle of resolution denotes the smallest visual angle (in arcminutes) at which an individual can distinguish two separate points, applicable prior to object recognition. In contrast, edge-to-edge critical spacing indicates the spatial range, extending beyond the target area, over which information is pooled for feature integration and object segmentation. Specifically, we compared the acuity thresholds or MAR between the unflanked stimulus condition and the edge-to-edge critical spacing threshold, which was derived only from flanked trials at the threshold stimulus size, across different eccentricities (figure 3*d*). A repeated-measures ANOVA revealed a significant interaction between eccentricity and measure type (E-E Spacing vs. MAR), F(3,15) = 8.82, *p* = 0.0013, *η*^2^ = 0.0035), indicating that the relationship between the two measures varied with eccentricity. Post hoc multiple comparisons confirmed that MAR and E-E spacing were comparable at the PLF (*p* = 0.41), but diverged at larger eccentricities (*p* = 0.018, at 10’, *p* = 0.002 at 15’ and *p* = 0.002 at 25’).

Perceptual mislocalizations, where the target’s position is confused with that of a flanker [56–58], often occur in crowding. In a four-alternative forced-choice task, there are three other possible options for an incorrect response, corresponding to a guess rate for mislocalizations of 33%. If there is an increased probability of responding to one of the flanker locations during incorrect trials, this suggests a bias toward that flanker location (an example for stimuli presented at the center of gaze and at 15’ eccentricity is shown in Figures 4*a* and *b*). On flanked trials, flankers were selected from a pool of four possible stimulus options with replacement, thus individual stimulus configurations did not necessarily include all four positions options and could contain duplicates. Since the upper and lower flanker locations were equidistant from the center of gaze, we next focused on the directional influence of the inner and outer flanker locations, where the inner flanker was positioned closer to the PLF and the outer flanker was slightly farther from it. Figure 4*c* shows the probability of reporting either the inner or outer flanker on incorrect trials. We observed a significant effect of eccentricity on mislocalization rates (repeated-measures ANOVA: F(3,15) = 5.07, p = 0.013, *η*^2^ = 0.005). Furthermore, there was a significant main effect of the type of mislocalization, indicating a difference between inner and outer mislocalizations (F(1,5) = 8.34, p = 0.034, *η*^2^ = 0.028); the inner flanker was more likely to be mislocalized as the target. However, the interaction between eccentricity and mislocalization type did not reach significance (F(3,15) = 1.82, p = 0.19, *η*^2^ = 0.01). Overall, these findings indicate that the effects of crowding are not uniform within the foveola. Specifically, the rate of change in critical spacing with eccentricity, quantified as the slope of the fitted function, was ∼ 0.04 within the foveola. This pattern aligns with the third possible outcome proposed in Figure 1*a*; crowding increases gradually within the foveola rather than remaining constant or showing a sharp discontinuity. Figure 5*a* shows the change in critical spacing (in arcminutes) as a function of stimulus eccentricity within the foveola. This prompts the question of how this increase compares with the magnitude of crowding effects observed at eccentricities outside the foveola. To address this question, we assessed the rate at which crowding increases from 1 to 6 deg of eccentricity in 4 of our subjects. Outside the boundaries of the foveola, the critical spacing was found to be larger, and the rate of change in critical spacing with eccentricity, quantified as the slope of the fitted function, was also higher (0.149). Figure 5*b* shows the relationship between critical spacing (in arcminutes) and eccentricities outside the foveola. Crucially, in the foveola, this rate is approximately 3.5 times smaller than extrafoveally, further highlighting the fact that the foveola is not only a non-uniform region, but crowding processes at this scale follow a different regime compared to extrafoveally.

**Figure 4:**
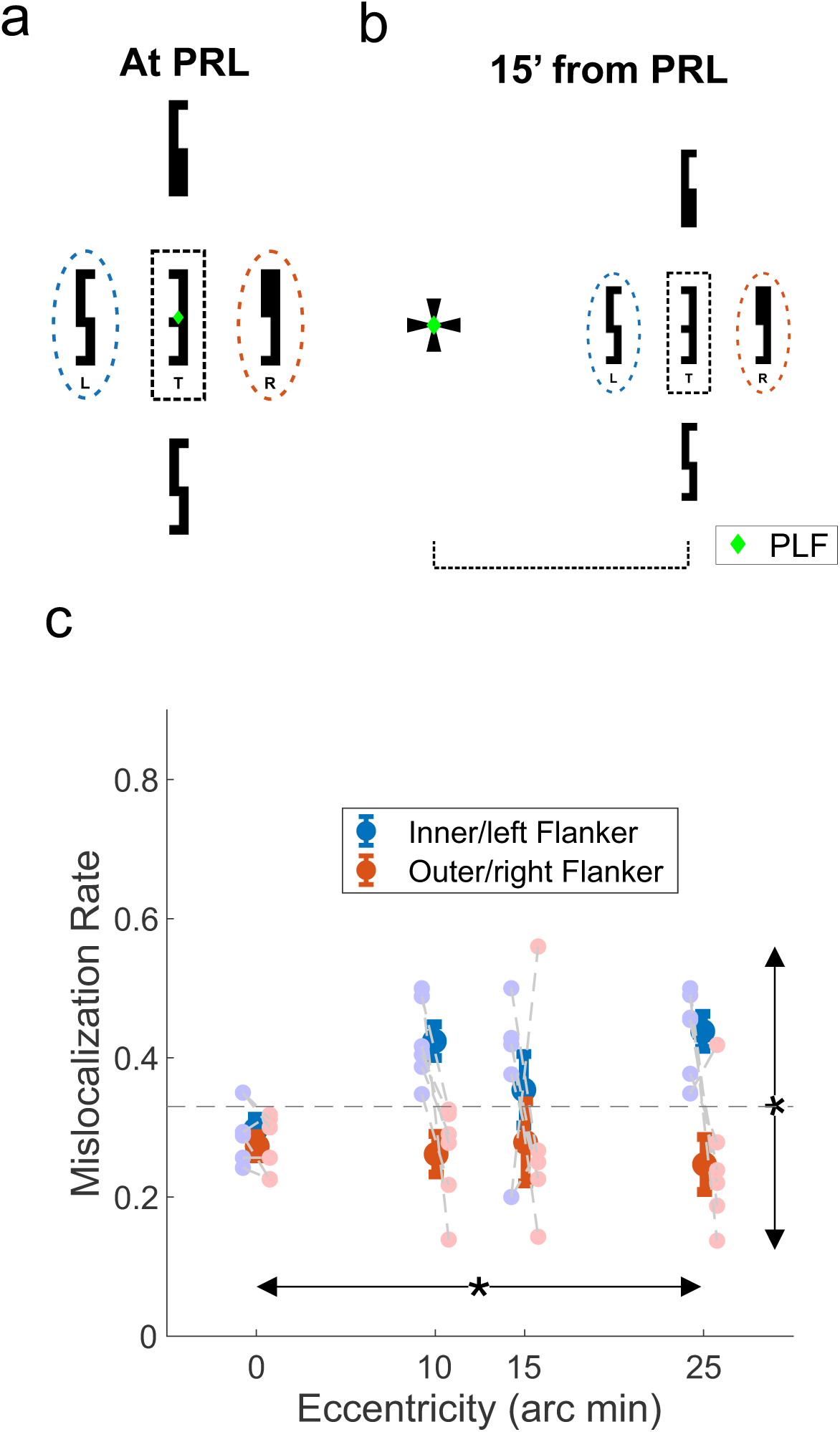
Mislocalization errors in the crowded condition. (a) Illustrates a crowded stimulus presented at the center of gaze (0’ eccentricity), with the target highlighted by a black dashed square. The leftward and rightward flankers are highlighted by blue and orange dashed ovals, respectively. At this eccentricity, the two flankers are equidistant from the preferred locus of fixation. (b) Depicts a crowded stimulus presented to the right of the center of gaze at 15′ eccentricity. Here the leftward/inner flanker is slightly closer to the center of gaze when compared to the rightward/outer flanker. (c) The probability of reporting either the inner/left or outer/right flanker as the target in incorrect trials is plotted for each stimulus eccentricity that was tested (repeated-measures ANOVA: p = 0.013 for eccentricity, p = 0.034). Error bars indicate s.e.m.. Asterisks denote significant main effects (^∗^*p <* 0.05, ^∗∗^*p <* 0.01).

**Figure 5:**
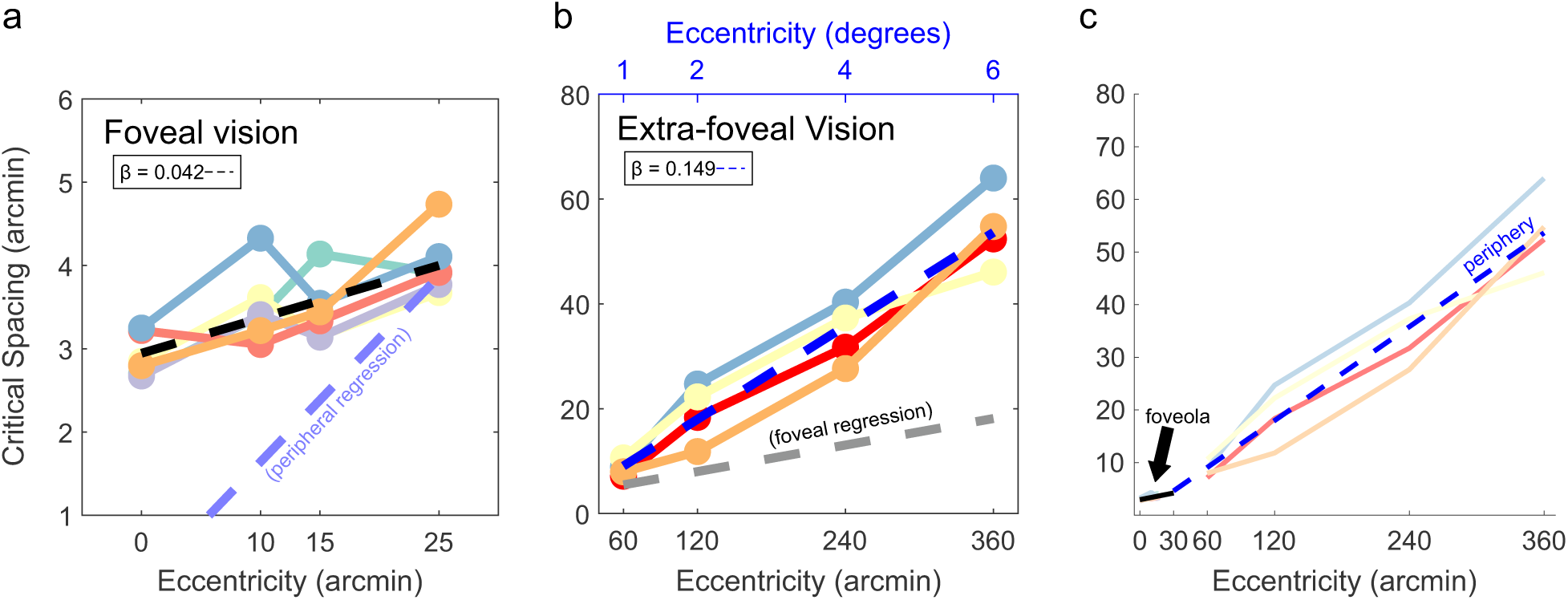
Foveal and extrafoveal crowding. (a) Center-to-center critical spacing as a function of eccentricity in the central fovea. The slope (*β*) of the fitted line (dashed black line) based on the average data is reported on top. Individual subjects are represented by different colors. The extrapolated fit based on the extrafoveal data is plotted as a reference. (b) Four of the same subjects measured in (a) tested at locations outside the central fovea. Conventions are the same as in (a). The extrapolated fit based on the foveal data is plotted as a reference. (c) Fits to the crowding growth functions show a markedly shallower foveal slope than extrafoveal.

**Figure 6:**
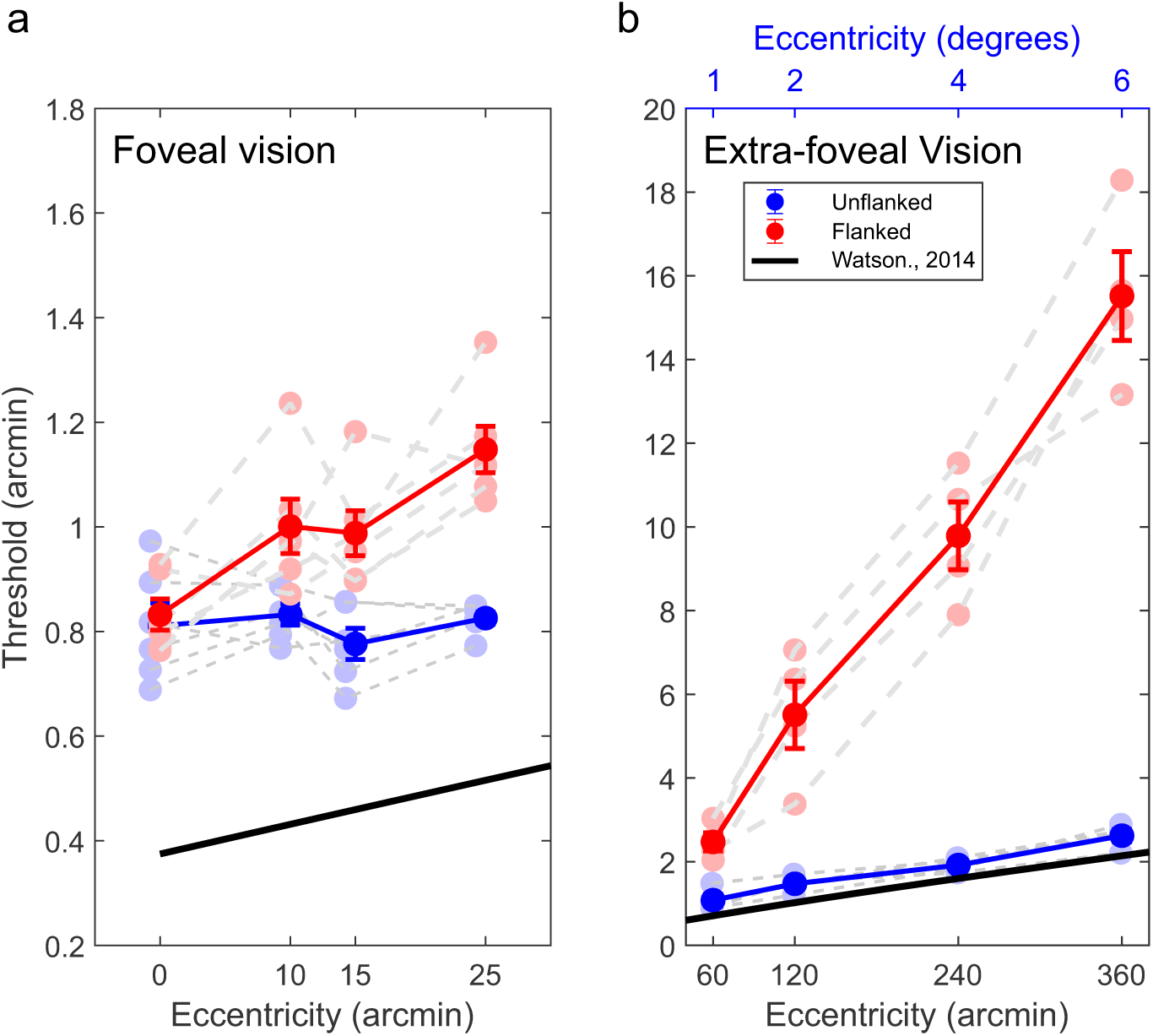
Crowding, acuity and midget retinal ganglion cell spacing foveally and extrafoveally. Edge-to-edge critical spacing and visual acuity thresholds for foveal (a) and extrafoveal (b) eccentricities. The solid black line in both panels represents the estimated midget retinal ganglion cell (mRGC) spacing (in arcminutes) based on Watson’s model (2014). Error bars indicate s.e.m.. While unflanked visual acuity closely matches mRGC spacing estimates across eccentricities, critical spacing increases more steeply. In the fovea, an offset from the mRGC limit may reflect the effects of habitual optical aberrations. Error bars are s.e.m..

Extrafoveally the phenomenon of visual crowding is generally characterized by the Bouma fraction, the ratio of critical spacing to eccentricity. Our results show that from 1 to 6 degrees the Bouma fraction remained relatively stable (1*^◦^*: 0.14 ± 0.02; 2*^◦^*: 0.16 ± 0.05; 4*^◦^*: 0.14 ± 0.02; 6*^◦^*: 0.15 ± 0.02) (see supplemental figure S2), consistent with work showing that the growth of crowding distance with eccentricity is supralinear but this supralinearity becomes obvious only at much larger eccentricities [59, 60]. Interestingly, the Bouma fraction from the current study was generally smaller than those reported in previous studies, where the Bouma fraction was observed to range between 0.4 and 0.5 [8, 58]. However, a few other studies have reported lower values, around 0.3 [9, 10, 60, 61], and even smaller values for tangential flanker orientation (ranging from 0.1 to 0.2) [1, 61] or when using different stimuli [62]. Choice of stimuli (4 out of 9 digits) in our study may have contributed to the lower Bouma’s factor. Crucially, within the foveola, the Bouma’s fraction varied with eccentricity (10’: 0.35 ± 0.04; 15’: 0.23 ± 0.02; 25’: 0.16 ± 0.02) (see supplemental figure S2), showing that foveal crowding cannot be described by a single Bouma’s fraction value. In the Foveola the Bouma fraction showed a linear decline with eccentricity, with a slope of –0.0118.

To determine whether the pattern of mislocalization observed within the foveola extends into the peripheral visual field, we next analyzed mislocalizations at greater eccentricities (eccentricities 1 to 6 degrees). At more peripheral locations (1*^◦^* – 6*^◦^*), mislocalization rates continued to increase with eccentricity (F(3,6) = 11.71, p = 0.0064, *η*^2^ = 0.0073). Although the main effect of mislocalization type (inner vs. outer) showed a strong trend (F(1,2) = 14.28, p = 0.063, *η*^2^ = 0.167), the interaction between eccentricity and mislocalization type was not significant (F(3,6) = 0.25, p = 0.86, *η*^2^ = 0.005). These results indicate that, as in the foveola, peripheral crowding is associated with increasing mislocalization errors with eccentricity, with a consistent directional bias.

To assess whether the observed acuity and crowding changes reflect retinal sampling limits, we compared our data to a model of midget RGC spacing. We used a theoretical model of the spacing between mRGC cells in the nasal retina, corresponding to the temporal visual field where stimuli were presented in the foveal and extra-foveal conditions [63]. Using the results shown in figure 3*d*, we determine whether the sampling limit of mRGC cells could account for the observed results. First, we compared the acuity measures from the unflanked condition across different eccentricities tested in both the fovea and extra-foveal locations. These comparisons are shown in 6*a*–*b*, with 6*a* presenting the foveal data (replicating the results from 3*d*) and 6*b* showing results from the extra-foveal condition. Similarly, we compared the edge-to-edge critical spacing measures across eccentricities with the changes in mRGC spacing. Overall, the slope relating mRGC spacing to unflanked visual acuity in the fovea was shallower than the model prediction (0.0001 ± 0.0042 vs. model = 0.0056; t(5) = –3.22, p = 0.023, d = 1.32; z = –1.99, p = 0.046), which may reflect the influence of habitual optical aberrations that were not corrected in this experiment. The slope for edge-to-edge critical spacing was, instead, steeper (0.0120 ± 0.0055; t(5) = 2.84, p = 0.036, d = 1.16; z = 2.20, p = 0.028). This indicates that mRGC spacing alone likely do not account for the changes in critical spacing with eccentricity. On the other hand, the extrafoveal data showed a closer match between unflanked acuity and mRGC spacing (periphery: mean = 0.0049 ± 0.0013 vs. model = 0.0062; t(3) = –1.93, p = 0.150, d = 0.96; z = –1.46, p = 0.144), likely because RGC density rather than optical aberrations is the limiting factor for acuity at these eccentricities [38]. However, the edge-to-edge critical spacing increased with eccentricity at a much steeper rate than predicted by mRGC spacing (0.0424 ± 0.0071 vs. model = 0.0062; t(3) = 10.23, p = 0.002, d = 5.11; z = 1.83, p = 0.068). While inter-RGC spacing predicts acuity in the extrafovea, it fails to account for crowding, consistent with the idea that crowding and acuity rest on different mechanisms. Note, however that prior work has shown that crowding zones subtend a nearly constant number of RGCs across 4 ° - 18.5 °[33], suggesting that the RGC layout may still provide an appropriate metric for characterizing crowding extrafoveally.

## Discussion

Here, we investigated the impact of visual crowding across the central 1*^◦^* foveola while precisely controlling for stimulus positioning at sub-foveal eccentricity by using high-precision eye tracking [52, 53]. Although this region is often studied as a single functional unit, our findings provide clear evidence that visual phenomena such as crowding vary systematically with eccentricity, even within this tiny part of the visual field. Crowding extent increased linearly with eccentricity in the foveola. Notably, the rate at which crowding magnitude increased with eccentricity was substantially slower than what has been reported extrafoveally, offering new insights into the distinctions between foveolar and extra-foveolar vision.

Although the existence of crowding in the central fovea has been debated, there is now compelling evidence that this region is not immune to crowding. Importantly this implies that Bouma’s law breaks at the center of gaze, raising the question of whether the effects of crowding are uniform in the central fovea, and if not, how the relationship between crowding and eccentricity at the foveal scale differs from the well-characterized increase in crowding with eccentricity in the extrafoveal visual field. Strasburger (2019) speculated that there may be a non-linear transition phase near the center of gaze (within 0.2*^◦^*), where critical spacing increases more gradually before conforming to a linear trend outside this region. Our findings shed light on this by revealing that the intensity of crowding is not uniform across the foveola; the rate of increase with eccentricity is approximately 3.5 times slower foveally than that observed from 1 to 6 deg outside this region (see figure 5*c*). Our data suggest that there are two phases in the growth of crowding with eccentricity, a slow phase within the boundaries of the foveola, and a fast phase, outside this region, supporting the idea that crowding follows a distinct scaling regime in the central fovea compared to more eccentric locations. A piecewise linear fit to group-level data revealed a break-point at 26.23 arcmin, with a shallow slope in the foveal region (slope = 0.042) and a steeper increase in the peripheral region (slope = 0.149).

Crowding, often considered a hindrance to visual discrimination, can be advantageous in natural viewing conditions. While it impairs the ability to distinguish individual elements by integrating information across pooling regions, this integration can also facilitate the perception of patterns in visual input [2, 64–66]. Gestalt mechanisms, such as grouping, mitigate interference from flanking objects when they are highly similar or aligned, allowing them to be perceived as a unified whole [67, 68]. This effect is particularly strong in natural scenes where multiple objects and textures form coherent structures. At the foveal scale, crowding mechanisms likely aid fine texture discrimination and shape grouping, supporting efficient processing of spatial redundancies. However, foveal crowding can hinder precise recognition tasks, such as when reading small text from a distance.

Our findings suggest that the underlying mechanism of crowding may be consistent across the visual field, but the rate at which its spatial extent grows differs between the foveola and periphery. This difference is likely driven by increased spatial integration outside the foveola, due to greater retinal ganglion cell (RGC) convergence, which leads to increased pooling of visual information [19, 24]. Several well-known factors differ between these two regions, including visual acuity [69–71], photoreceptor packing [24], blood supply [72] and ganglion cell mapping [73]. Previous research also supports the existence of distinct functional channels (*e.g.* color, motion), which are present throughout the retina, though their relative contribution may differ across regions due to anatomical variations [73, 74]. It is possible that the different rate of change of the crowding extent with eccentricity in the central fovea vs extrafoveally may arise from the larger receptive field pooling and loss of 1-to-1 photoreceptor to ganglion cell connections beyond the 1-deg fovea [33], leading to broader spatial integration and increased interference from neighboring elements. In fact, although previous work suggests that the private line extends beyond he central fovea [19], work from Rossi and Roorda (2010) suggests that it may be limited to a much smaller area as acuity under diffraction limited conditions matches with cone density only within the central 1 deg fovea but not outside.

Beyond retinal-level explanations, recent work has emphasized the role of cortical architecture in shaping crowding. The observed increase in critical spacing with eccentricity has been attributed to a reduction in cortical distance between the stimuli representations, reflecting how information is organized in the brain [5, 34, 75]. It has been proposed that the product of the Bouma’s function (*i.e.*, radial crowding distance increases linearly with eccentricity) and the cortical magnification function (*i.e.* radial cortical magnification decreases inversely with eccentricity) is approximately constant [5], leading to conservation across eccentricity of the threshold critical spacing on the cortical surface. Consistent with this Coates et al. [34] demonstrated that in the parafovea the spatial extent of crowding remains stable when expressed in cortical units. This suggests that cortical distance, shaped by cortical magnification, plays a key role in determining crowding effects at least extrafoveally. Importantly, recent work has highlighted how the critical cortical crowding distance may not be constant, as initially assumed, but may increase rapidly within the fovea and reach an asymptote only beyond ≈ 5 deg [76]. Yet, relatively little is known about how visual functions and the cortical magnification gradient are modulated within the central fovea. High-resolution fMRI studies [36, 77], have shown that there is a marked cortical magnification gradient within the foveola.

In conclusion, our study demonstrates that visual crowding exhibits a systematic gradient within the central foveola, revealing that even minute increases in eccentricity lead to measurable changes in critical spacing. Interestingly, the rate at which crowding extent increases with eccentricity is 3.5 times slower within the central fovea. These differences are likely due to differences both in cortical magnification and retinal convergence between foveal and extrafoveal representations. The presence of a crowding gradient within the central foveola is consistent with the idea that cortical magnification is characterized by a gradient even at this scale [36, 77]. On the other hand, the clear dichotomy in the rate of increase between the foveal and extrafoveal crowding may reflect the fact that the region fully characterized by a private-line connection between cones and RGCs extends only to the central 1 deg region, in line with previous work by Rossi and Roorda (2010). Ultimately, these results show that the central foveal vision is less uniform than previously assumed and is not merely an extension of the processing principles and regimes that apply to the rest of the visual field.

## Methods and Procedures

### Observers

6 participants with normal vision took part, including 5 naive individuals and one experienced observer who is also one of the authors took part in the study. The group comprised 2 males and 4 females, aged between 18 and 29 years. During the initial screening, all participants demonstrated at least 20/20 Snellen acuity with or without the need for corrective lenses. Ethical approval for this research study was obtained from the University of Rochester’s Research Subjects Review Board. Subjects attended an initial screening session, which involved a comprehensive explanation of the experiment and a thorough review of the consent form materials. After understanding the information and voluntarily agreeing to participate, informed consent was obtained and documented.

### Stimuli and Apparatus

Stimuli were presented monocularly to the right eye while the left eye was covered. Eye movements were recorded with high precision using a custom-made digital Dual Purkinje Image (dDPI) eye tracker [52], operating at a sampling rate of 340Hz. This system had minimal internal noise (below 1 arcminute) and a spatial resolution of at least 1 arcminute [52, 78, 79]. To minimize noise and enhance eye-tracking precision, the observer’s head was immobilized with a dental-imprint bite bar and head-holder. The stimuli were displayed on an ASUS PG258Q LCD monitor with a vertical refresh rate of 200Hz and a spatial resolution of 1920 × 1080 pixels. The monitor was positioned at either 3 or 5 meters from the observer, corresponding to pixel sizes of 0.25 arcminutes and 0.19 arcminutes, respectively.

Stimuli consisted of digits (3, 5, 6, and 9) from the Pelli number-font [12] and were presented individually at the center of the display for 500 ms. The Pelli number font is specifically designed to test crowding within the fovea. Its vertically elongated design (aspect ratio 5:1) allows for a smaller center-to-center separation between the target and flanker compared to fonts with equal aspect ratios, thus providing a robust estimate of crowding thresholds. It has further been shown that the critical spacing estimates using the font are independent from the spacing-to-size ratio within the fovea [12], also reconfirmed by our previous work [17]. On flanked trials, flankers were drawn from the same pool of four possible digits, selected with replacement. As a result, not all four flanker identities necessarily appeared in a given configuration, and some flankers could be duplicated. The stimuli were presented at maximum contrast in black text on an uniform gray background and were controlled using EyeRIS [53], a custom-developed system enabling gaze-contingent display control. Stimuli were presented either at the center of the display (0’ eccentricity) or at one of three other eccentricities to the right or left of the center of gaze (10’, 15’, 25’). On trials where stimuli appeared at the center of the display (0’ eccentricity), in addition to the stimulus we presented four peripheral arches centered on the target to help subjects maintain their gaze near the center. On the other hand, when stimuli were presented away from the center of gaze, a 10×10 arcminute fixation marker appeared at the center of the display to facilitate precise fixation behavior. When flankers were present, the separation between the edge of the target and the flanker (flanker spacing) was set to be 1.4 times the width of the stimulus (spacing-to-size ratio or nominal spacing). To compute the corresponding edge-to-edge (E–E) spacing, we first determined the center-to-center spacing between the target and flanker at threshold, and then subtracted half the target width and half the flanker width (*i.e.*, the stimulus width), yielding the true E–E spacing.

### Experimental Paradigm

Data collection involved multiple experimental sessions, each lasting about one hour, with each subject completing an average of 5 sessions. Each session commenced with setup operations to ensure optimal observer positioning and eye tracker calibration. A two-step gaze-contingent calibration procedure was performed to map the eye tracker’s output to visual angles, enhancing the localization of the preferred retinal locus of fixation. During the first phase (automatic calibration), observers sequentially fixated on a 3×3 grid, while the second phase (manual calibration) allowed observers to confirm or refine the calibration by fixating on the same grid points while manually adjusting a gaze contingent marker to overlap with their gaze direction at each location. Before each trial, manual calibration was repeated for the central position to compensate for possible head movements.

Each trial began with a 10×10 arcminute fixation point at the screen center, followed by a 400 ms delay to prevent aftereffects from the fixation point. The target stimulus was then presented, and participants identified it by selecting one of four possible digits using a remote controller. Fixation trials, in which participants maintained fixation on a 10×10 arcminute marker at the display center, were interspersed approximately every 30-50 task trials, with subjects instructed to maintain fixation for 2-5 seconds. Target acuity and crowding threshold (critical spacing) was determined using the Parametric Estimation by Sequential Testing (PEST) procedure [55], where both the target size and flanker spacing changed based on subject’s performance which was set to converge at 62.5% for a 4AFC task.

All subjects started the initial experimental session with the stimuli presented at the center of gaze in the uncrowded condition. After the first 100 trial block, condition blocks where then randomized by eccentricity (either presented at 0, 10, 15, or 25 eccentricity temporally) and crowding condition (unflanked or crowded). All subjects were run on each unique block type at least twice to ensure repeatable thresholds estimates. A subset of subjects was also tested in the nasal visual field; however, because no systematic differences were observed between nasal and temporal thresholds, we restricted our analysis to the temporal dataset.

### Refractive Correction

In the current study, we employed a badal lens setup to compensate for spherical refractive error. This device allows participants to make precise adjustments, correcting for different amounts of spherical defocus and achieving high acuity vision. The Badal lens modifies the effective focal length of the lens system, ensuring that all stimuli are presented at the same size in visual angle by accounting for the introduced magnification factor.

To guide the correction process, we used the Atchison’s model [80] for our Badal lens. This model describes a relay system that helps achieve accurate corrections with limited distortions. Participants utilized this guidance to correct for varying amounts of spherical defocus and achieve corrections to 20/20 vision or better. Additionally, to assist participants in their subjective correction, we integrated a duo-chrome test with a red and green screen. Subjects ensured that the target was equally clear on both the red and green backgrounds before proceeding. This measure facilitated the fine-tuning of visual adjustments, contributing to the accuracy of participants’ corrective experiences throughout the study.

### Data Analysis

Eye Movements: Eye movements were categorized as saccades (including microsaccades) and ocular drift. Automatic classification was followed by a manual review by an expert experimenter. Trials with saccades, blinks, or poor tracking during stimulus presentation were excluded. On the other hand, when stimuli were presented away from the center of gaze (eccentricities 10, 15 and 25 arcminutes), trials where the gaze position deviated more than 30 arcminutes from the central fixation point were discarded. Ocular drift was examined during fixation periods far from saccades or blinks, revealing stationary behavior with consistent speed. On average, 32.8% ± 23.7% of trials were discarded across subjects based on these criteria. The same exclusion was used for the larger eccentricities tested (1 to 6 degrees).

Estimation of Acuity Thresholds: Visual acuity was calculated both as units of stimulus width (arcminutes) and as minimum angle of resolution (MAR). Visual acuity thresholds, representing the minimum stimulus width for reliable performance above chance (62.5% correct, with a 25% chance level), were determined using a cumulative Gaussian psychometric function [81]. Lower thresholds indicated better acuity. To convert Pelli digits’ strokewidth to MAR, instead of taking 1/5 of the stimulus width as with the tumbling E, MAR was defined as 1/2 of the stimulus strokewidth [12]. Therefore, an optotype that was 2′ wide would correspond to the 20/20 Snellen MAR line. Individuals psychometric fits are shown in supplemental figure S3.

### Analysis of performance

Statistical comparisons were performed using a two-way ANOVA with subject and condition as factors, followed by Tukey-Kramer post hoc tests implemented in MATLAB. Effect sizes were quantified using eta squared (*η*^2^) for each main effect and interaction. Pairwise comparisons between conditions were assessed using paired two-tailed t-tests, and each was accompanied by Bayes factor estimation (*BF*_10_), using the toolbox (Github repository), and Cohen’s d to quantify the strength and magnitude of the observed effects.

All data and MATLAB scripts used to create the figures in the manuscript have been uploaded onto the Open Science Framework repository.

## ACKNOWLEDGMENTS

This work was supported by the National Institutes of Health grants R01EY029788-01 (to M.P.) and by and NIH EY001319 grant to the Center for Visual Science at the University of Rochester.

## Supplementary Figures

**Figure S1:**
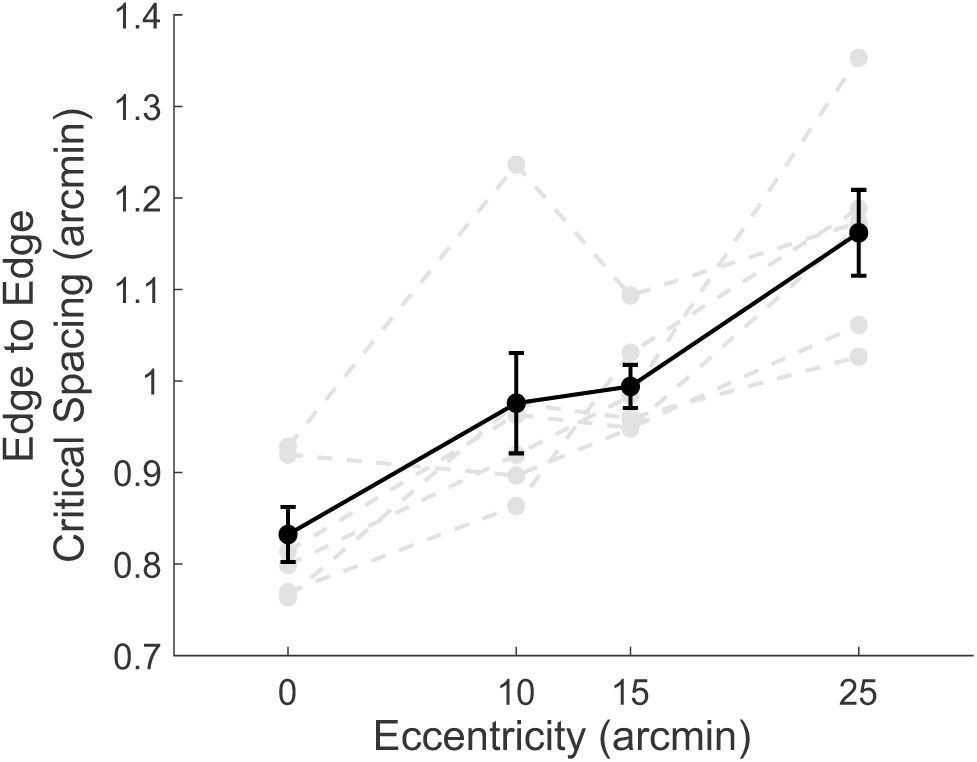
Edge-to-Edge spacing thresholds in the foveola. Edge-to-Edge spacing thresholds, represented as the distance between the inner edge of the target and the inner edge of the flanker for crowded stimuli, are plotted as a function of eccentricity. Error bars indicate s.e.m..

**Figure S2:**
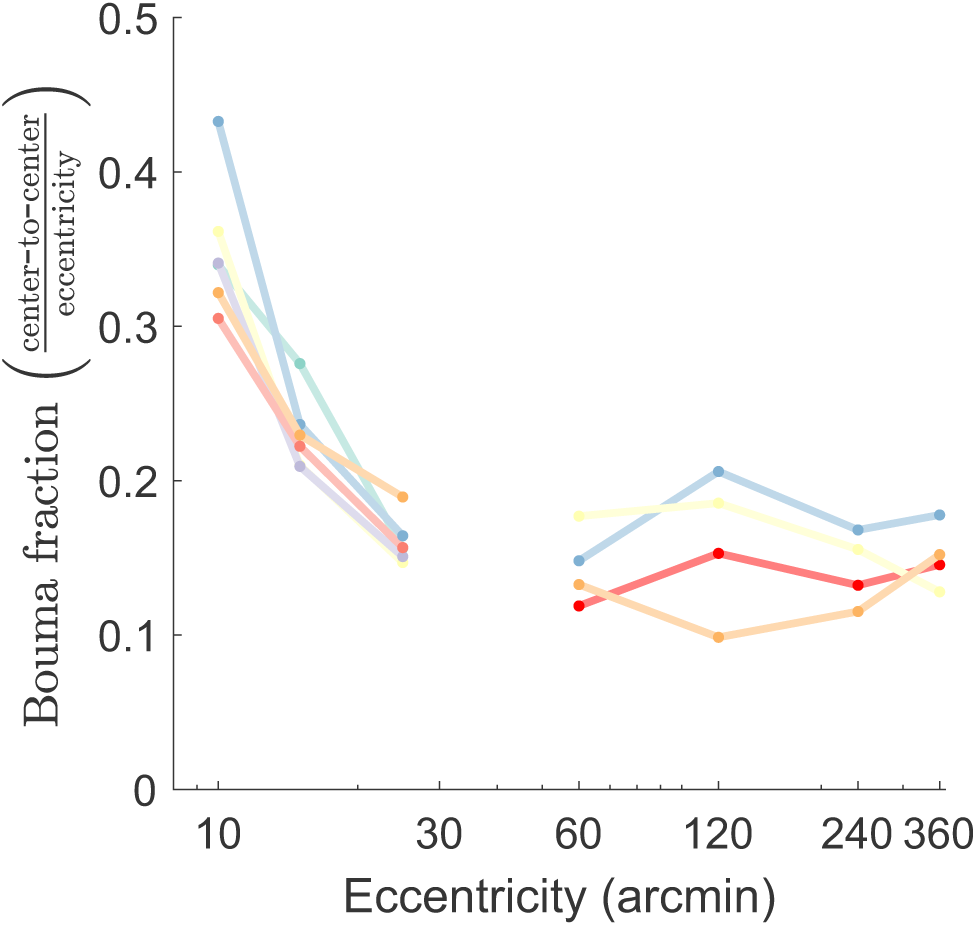
Bouma fraction across eccentricities. Colored lines represent individual subject data for both foveolar and extrafoveal conditions. The average trend across subjects shows a systematic decline in Bouma fraction within the foveola (slope of –0.0118), while the extrafoveal fit remains stable with a near-zero slope.

**Figure S3:**
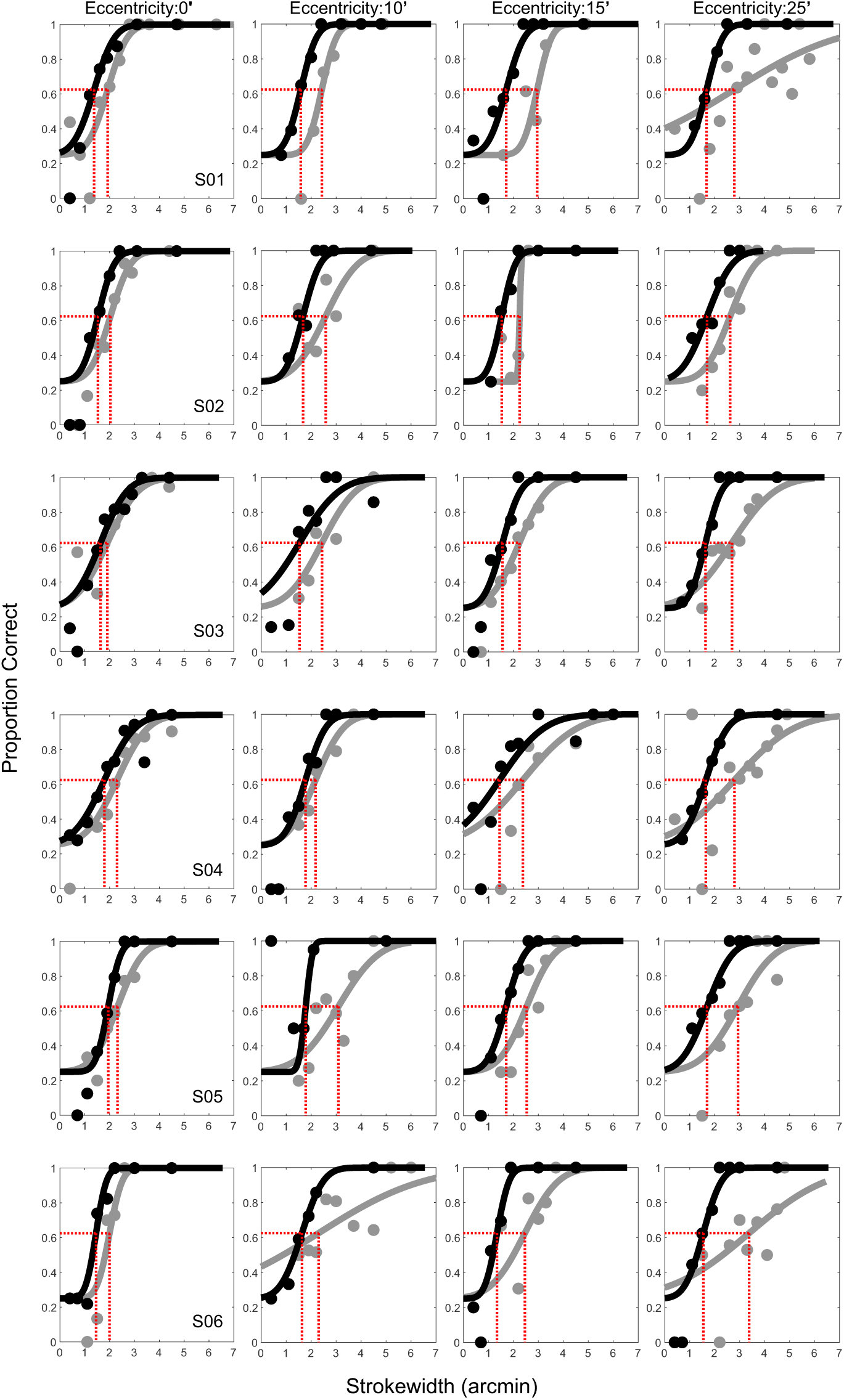
Psychometric functions for individual observers. Performance is plotted as a function of stimulus stroke width (in arcminutes) for the unflanked (black) and flanked (gray) conditions. Individual subjects are represented as separate rows, while each column represents the different eccentricities tested.

